## Supplementary material for "Mosaic cis-regulatory evolution drives transcriptional partitioning of HERVH endogenous retrovirus in the human embryo": supp. file 4

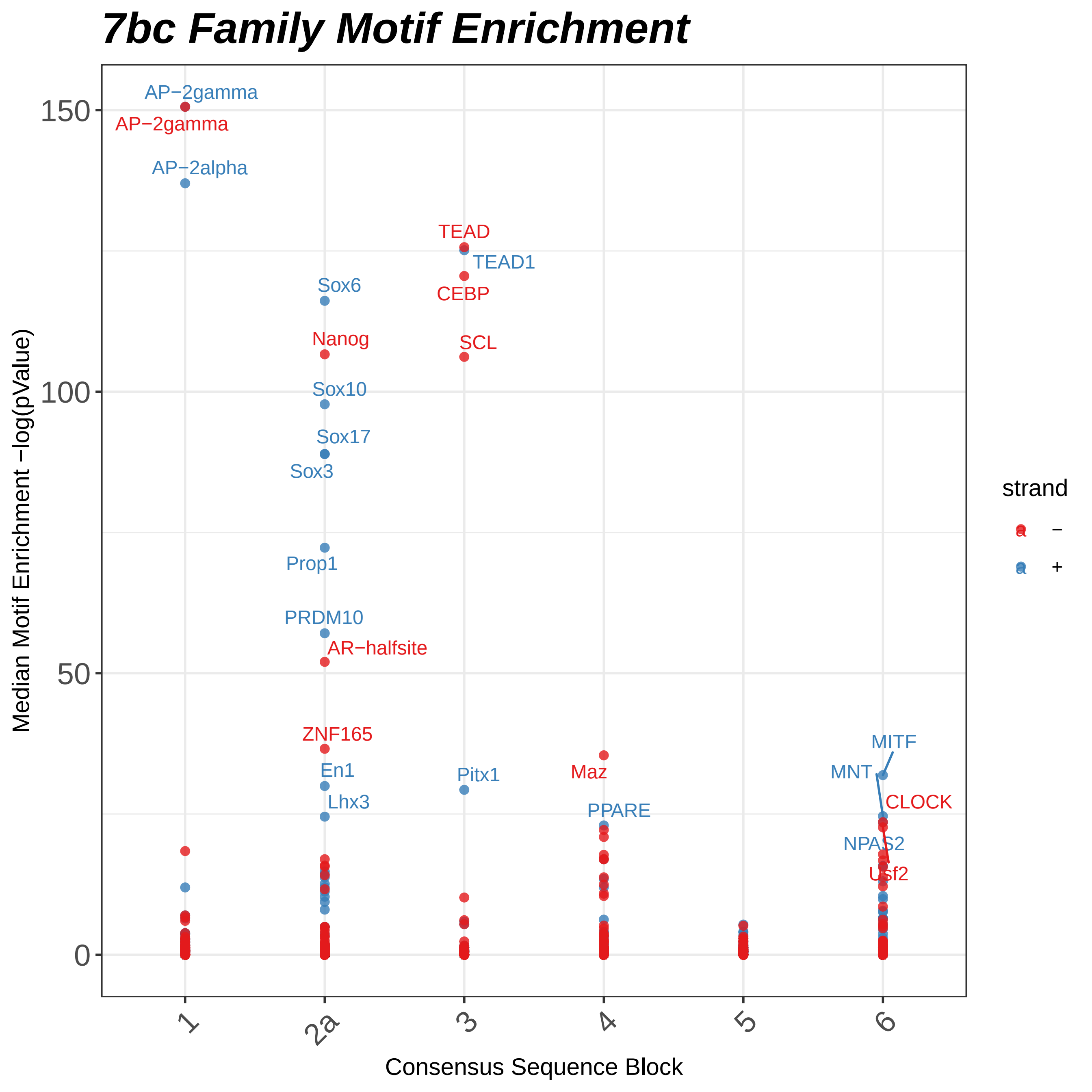
7bc Family Motif Enrichment
AP−2gamma
AP−2gamma
AP−2alpha
TEAD
TEAD1
CEBP
SCL
Sox6
Nanog
Sox10
Sox17
Sox3
Prop1
PRDM10
AR−halfsite
ZNF165
En1
Lhx3
Pitx1
Maz
PPARE
MITF
MNT
CLOCK
NPAS2
Usf2
150
100
50
0
Median Motif Enrichment −log(pValue)
1
2a
3
4
5
6
Consensus Sequence Block
strand
−
+

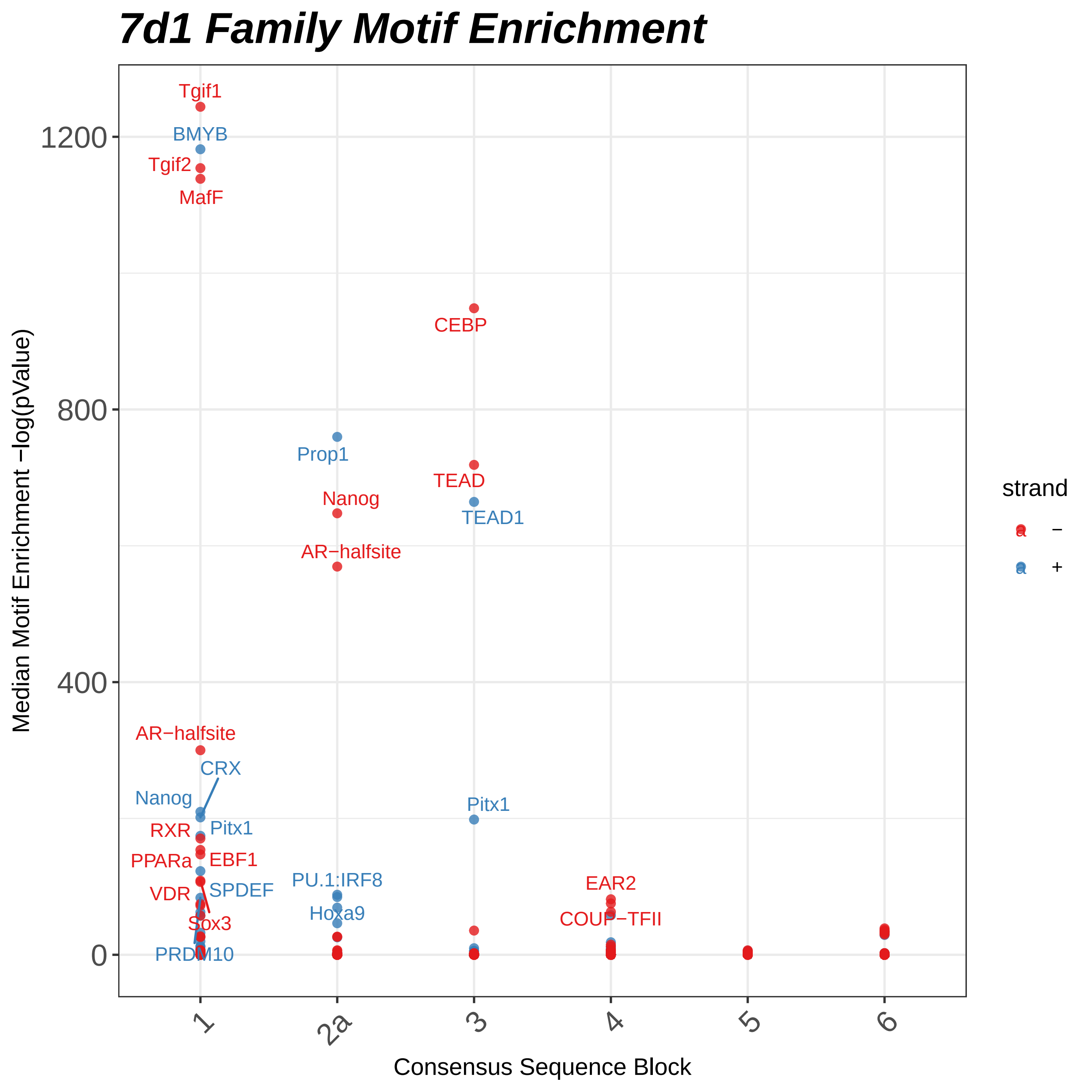

7d1 Family Motif Enrichment
Median Motif Enrichment −log(pValue)
Consensus Sequence Block
1200
800
400
0
1
2a
3
4
5
6
strand
−
+
Tgif1
BMYB
Tgif2
MafF
CEBP
Prop1
TEAD
Nanog
TEAD1
AR−halfsite
AR−halfsite
CRX
Nanog
Pitx1
RXR
Pitx1
PPARa
EBF1
VDR
SPDEF
PU.1:IRF8
EAR2
Hoxa9
Sox3
COUP−TFII
PRDM10

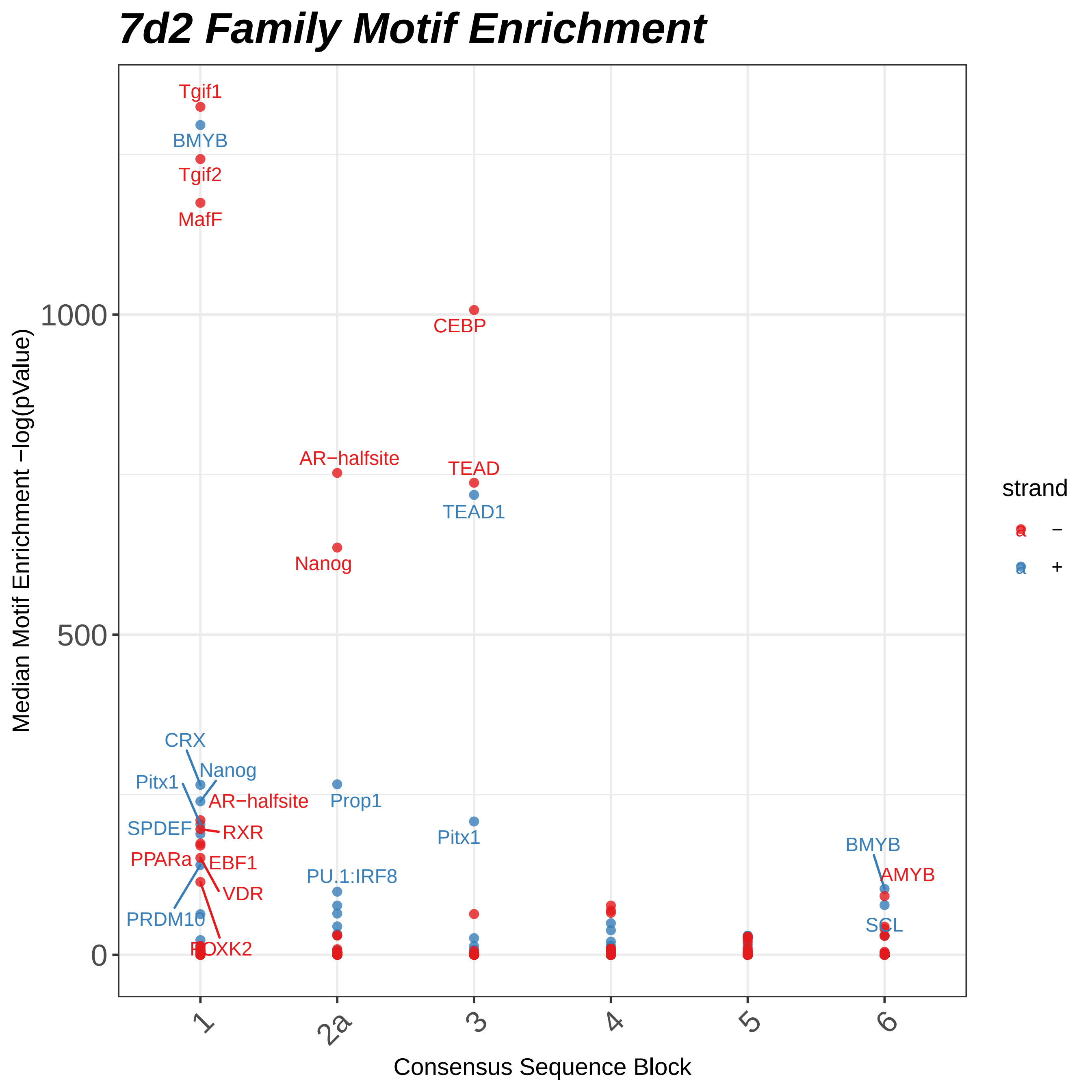

7d2 Family Motif Enrichment
Median Motif Enrichment −log(pValue)
Consensus Sequence Block
1000
500
0
1
2a
3
4
5
6
strand
−
+
Tgif1
BMYB
Tgif2
MafF
CEBP
AR−halfsite
TEAD
TEAD1
Nanog
CRX
Nanog
Pitx1
AR−halfsite
Prop1
SPDEF
RXR
Pitx1
PPARa
EBF1
BMYB
AMYB
PU.1:IRF8
VDR
PRDM10
SCL
FOXK2

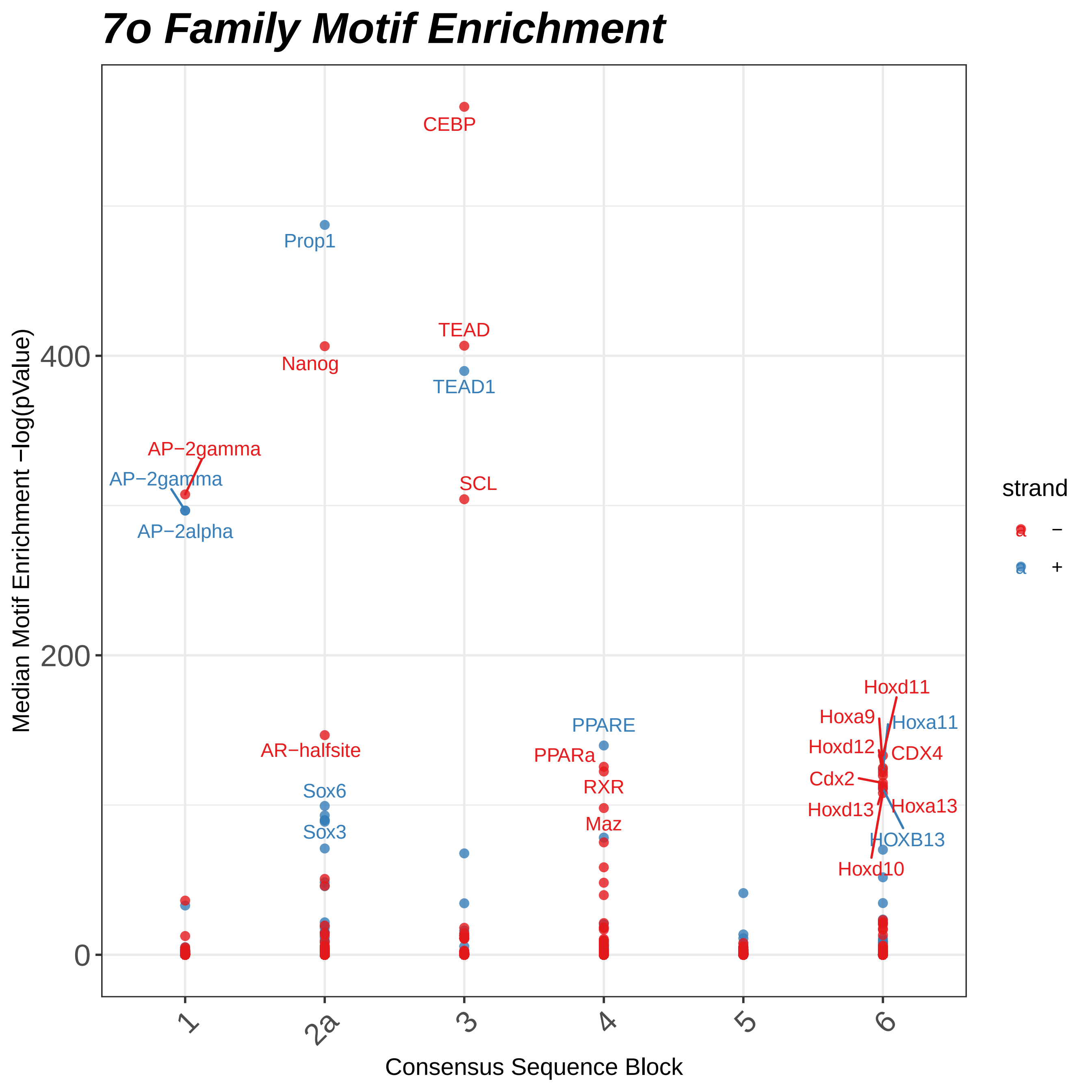
7o Family Motif Enrichment
Median Motif Enrichment −log(pValue)
0
200
400
CEBP
Prop1
TEAD
Nanog
TEAD1
AP−2gamma
AP−2gamma
SCL
AP−2alpha
Hoxd11
Hoxa9
Hoxa11
PPARE
Hoxd12
CDX4
AR−halfsite
PPARa
Cdx2
RXR
Sox6
Hoxd13
Hoxa13
Maz
Sox3
HOXB13
Hoxd10
strand
−
+
1
2a
3
4
5
6
Consensus Sequence Block

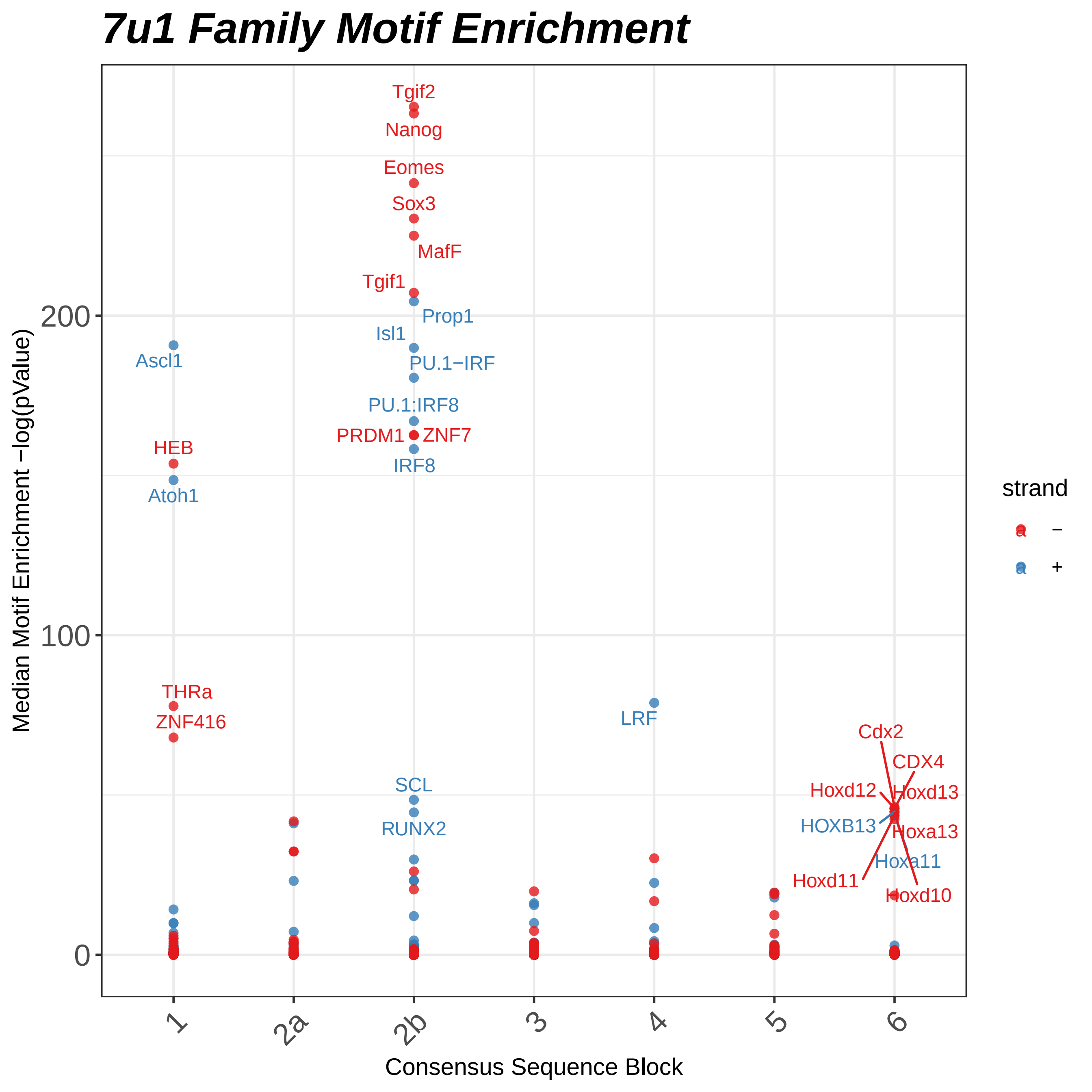
7u1 Family Motif Enrichment
Median Motif Enrichment −log(pValue)
Consensus Sequence Block
200
100
0
1
2a
2b
3
4
5
6
strand
−
+
Tgif2
Nanog
Eomes
Sox3
MafF
Tgif1
Prop1
Isl1
PU.1−IRF
PU.1:IRF8
PRDM1
ZNF7
IRF8
Ascl1
HEB
Atoh1
THRa
ZNF416
LRF
SCL
RUNX2
Cdx2
CDX4
Hoxd12
Hoxd13
HOXB13
Hoxa13
Hoxa11
Hoxd11
Hoxd10

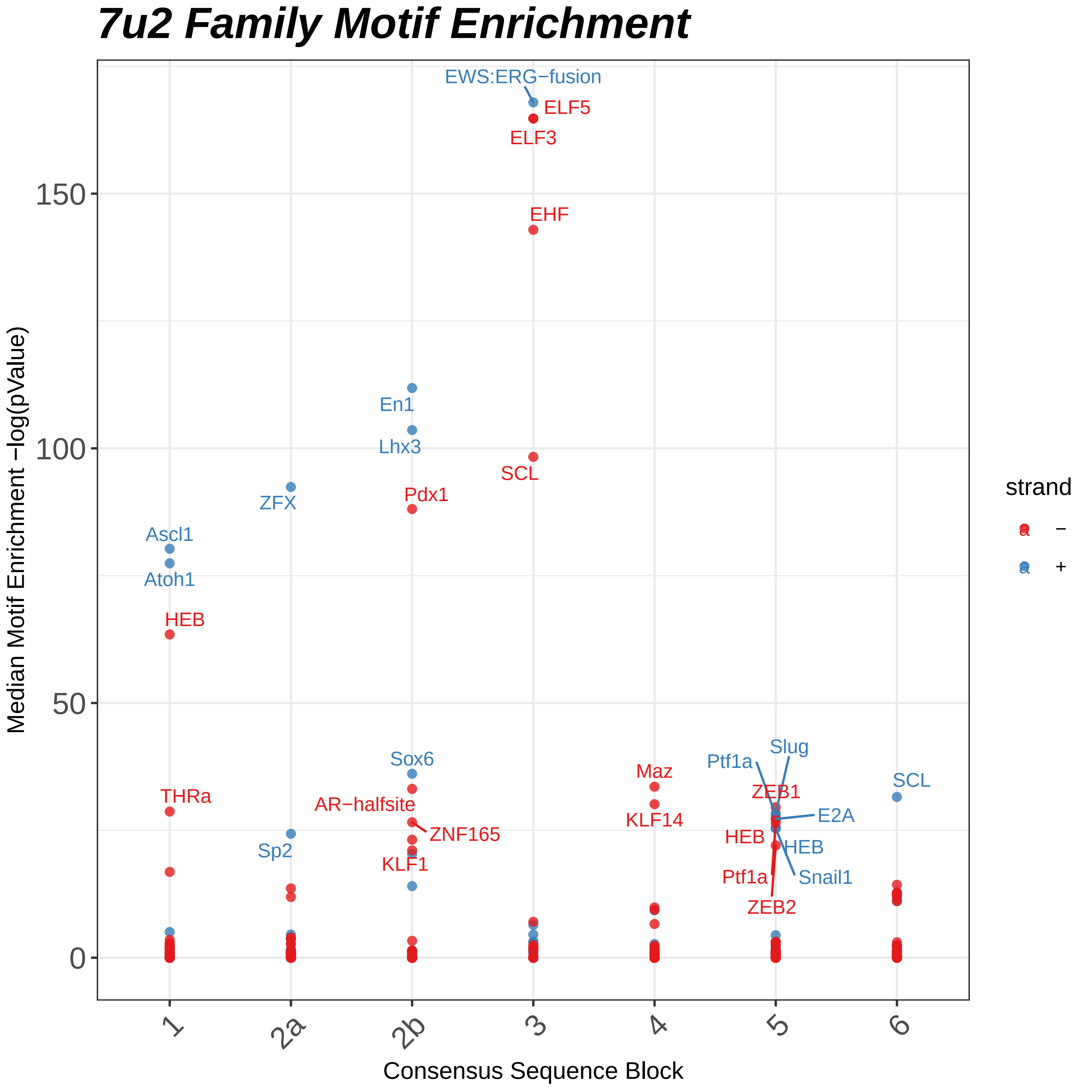
7u2 Family Motif Enrichment
EWS:ERG−fusion
ELF5
ELF3
EHF
En1
Lhx3
SCL
ZFX
Pdx1
Ascl1
Atoh1
HEB
Sox6
Maz
Slug
Ptf1a
ZEB1
SCL
THRa
AR−halfsite
E2A
KLF14
ZNF165
HEB
HEB
Sp2
KLF1
Ptf1a
Snail1
ZEB2
150
100
50
0
1
2a
2b
3
4
5
6
Median Motif Enrichment −log(pValue)
Consensus Sequence Block
strand
−
+

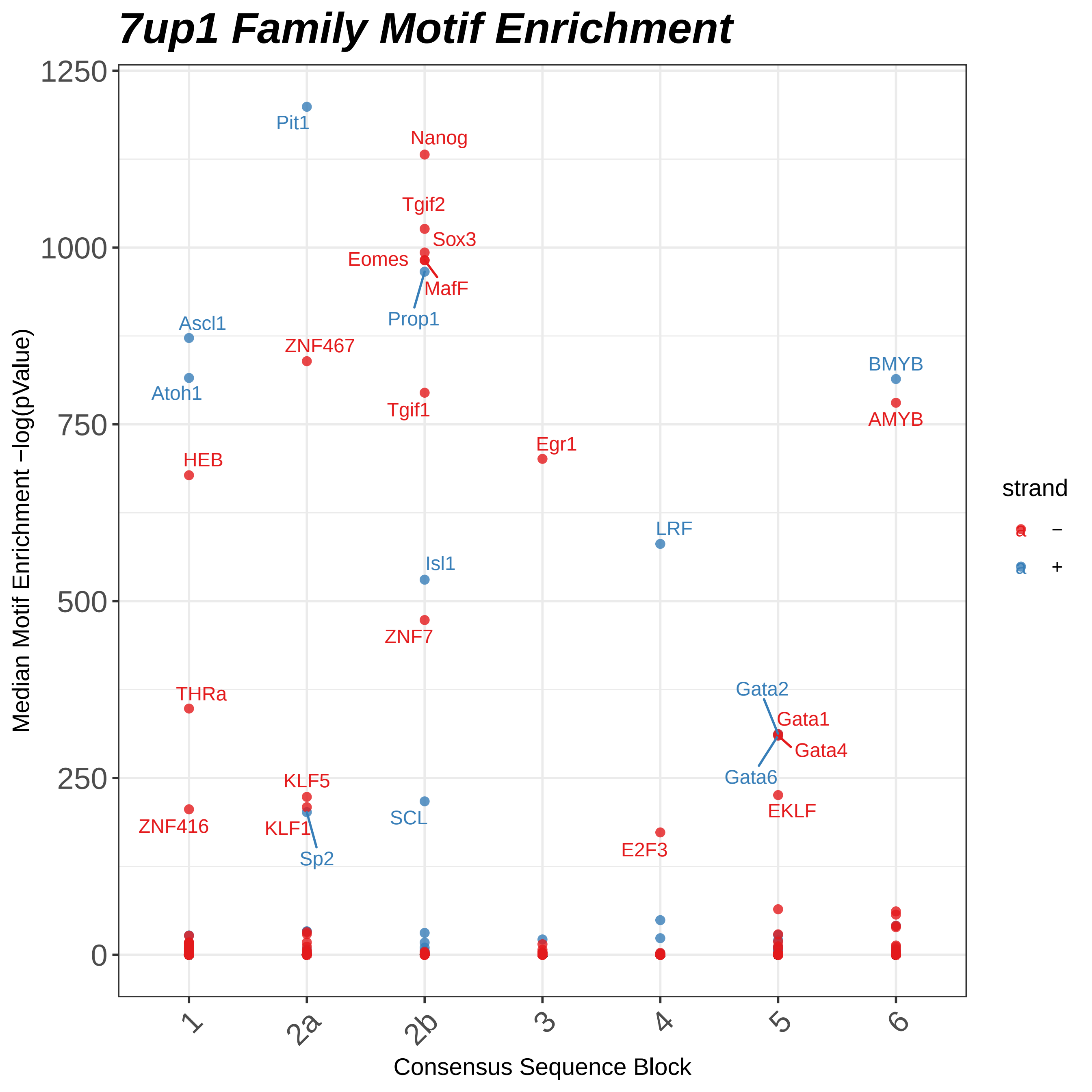
7up1 Family Motif Enrichment
Median Motif Enrichment −log(pValue)
1250
1000
750
500
250
0
Consensus Sequence Block
1
2a
2b
3
4
5
6
strand
−
+
Pit1
Nanog
Tgif2
Sox3
Eomes
MafF
Prop1
Ascl1
ZNF467
Atoh1
Tgif1
BMYB
AMYB
HEB
Egr1
LRF
Isl1
ZNF7
THRa
Gata2
Gata1
Gata4
Gata6
KLF5
EKLF
ZNF416
KLF1
SCL
Sp2
E2F3

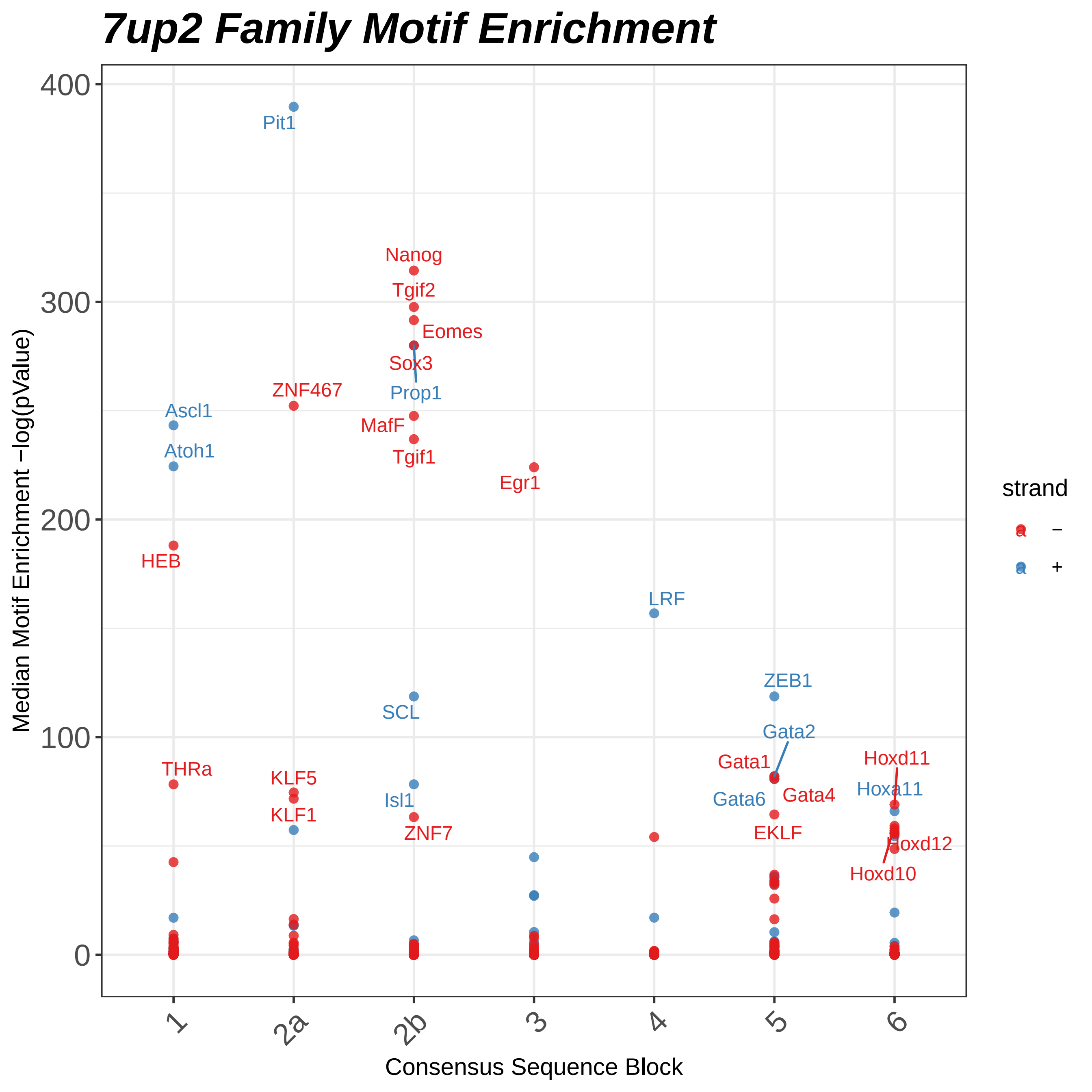
7up2 Family Motif Enrichment
Median Motif Enrichment −log(pValue)
400
300
200
100
0
Pit1
Nanog
Tgif2
Eomes
Sox3
Prop1
ZNF467
Ascl1
MafF
Atoh1
Tgif1
Egr1
HEB
LRF
ZEB1
SCL
Gata2
Gata1
Hoxd11
THRa
KLF5
Hoxa11
Isl1
Gata6
Gata4
KLF1
ZNF7
EKLF
Hoxd12
Hoxd10
1
2a
2b
3
4
5
6
Consensus Sequence Block
strand
–
+

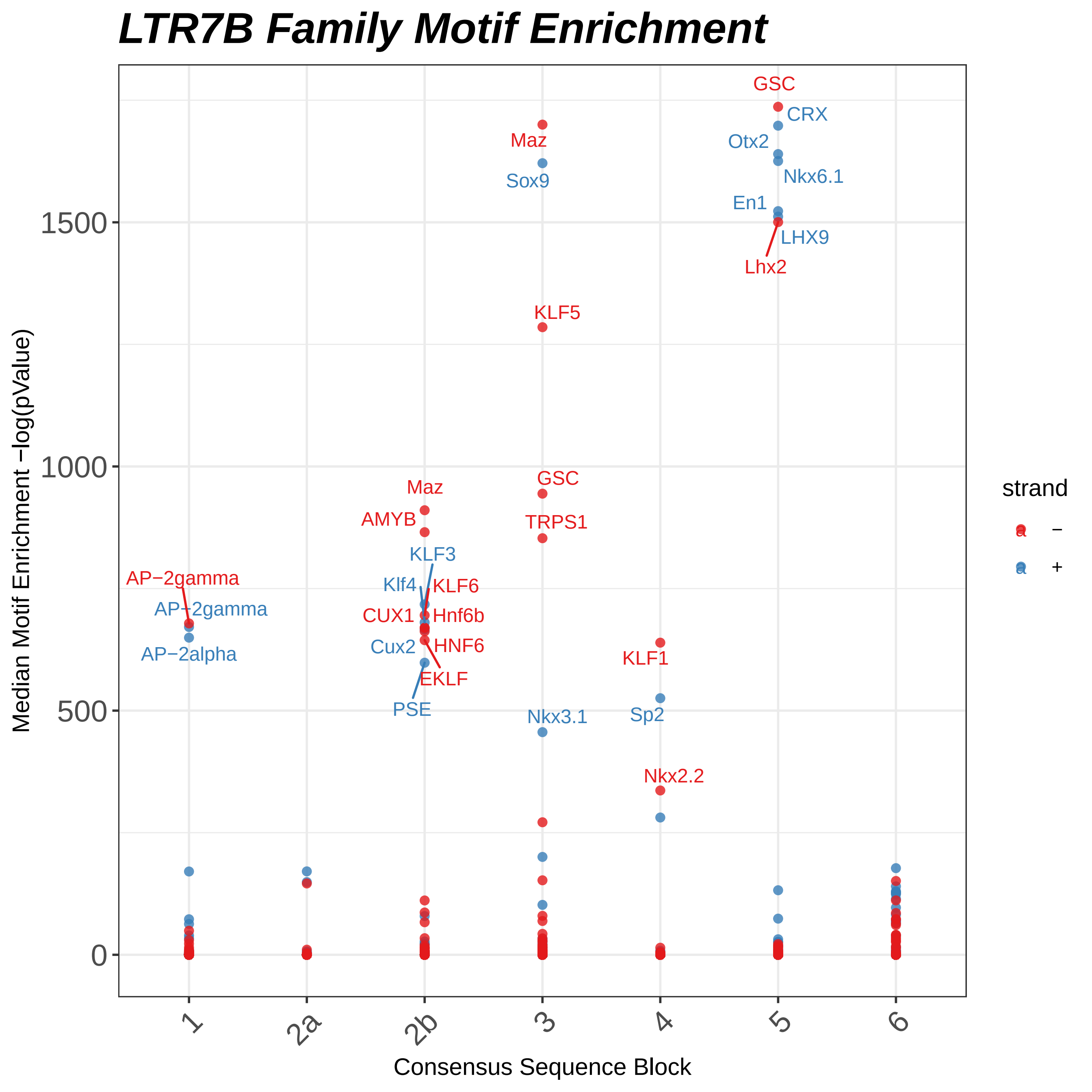
LTR7B Family Motif Enrichment
Median Motif Enrichment −log(pValue)
Consensus Sequence Block
1500
1000
500
0
1
2a
2b
3
4
5
6
strand
−
+
GSC
CRX
Otx2
Nkx6.1
En1
LHX9
Lhx2
Maz
Sox9
KLF5
GSC
TRPS1
Maz
AMYB
KLF3
Klf4
KLF6
CUX1
Hnf6b
Cux2
HNF6
EKLF
PSE
AP−2gamma
AP−2gamma
AP−2alpha
KLF1
Sp2
Nkx3.1
Nkx2.2

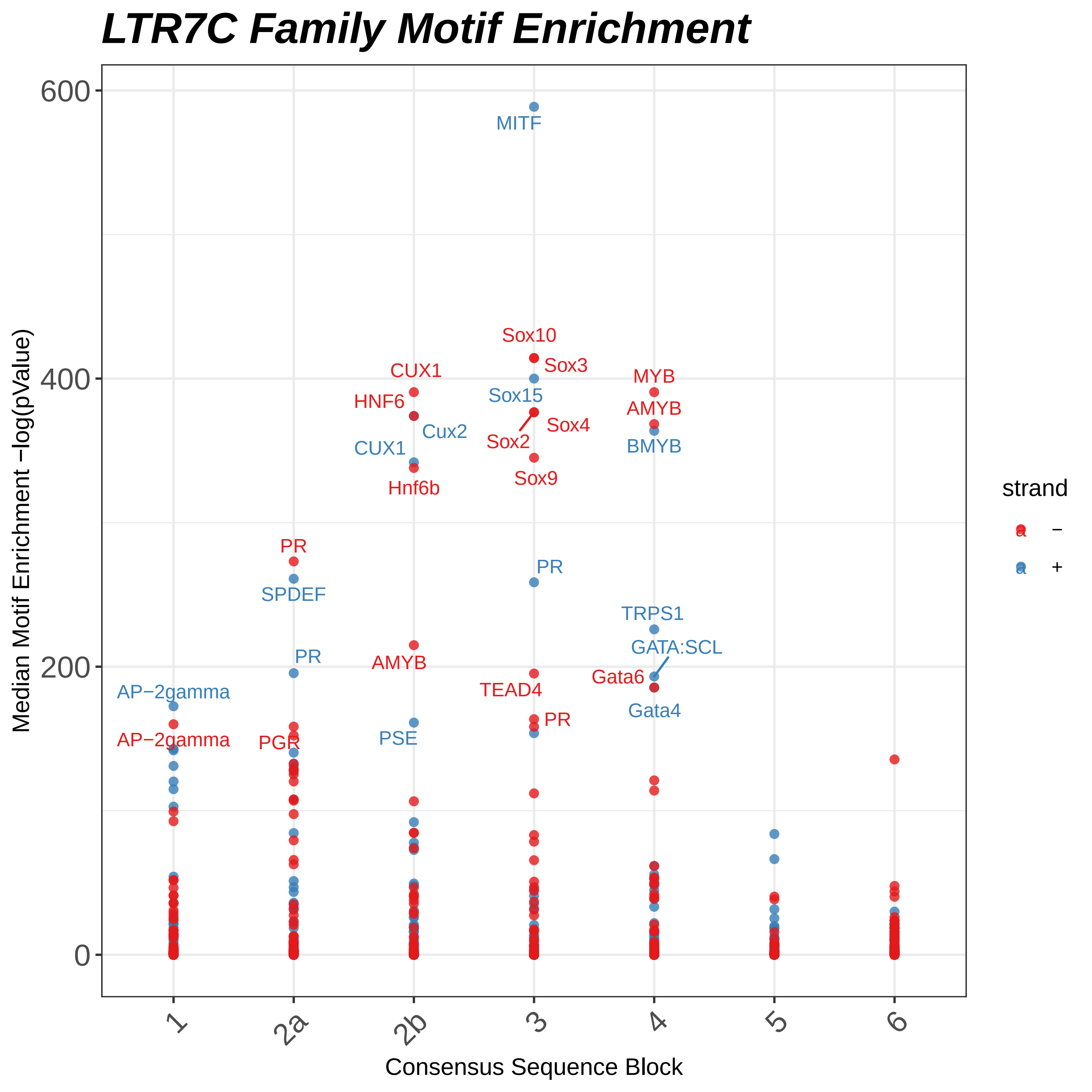

LTR7C Family Motif Enrichment
600
400
200
0
Median Motif Enrichment −log(pValue)
MITF
Sox10
Sox3
Sox15
Sox4
Sox2
Sox9
CUX1
HNF6
Cux2
CUX1
Hnf6b
MYB
AMYB
BMYB
PR
SPDEF
PR
PR
AMYB
TEAD4
TRPS1
GATA:SCL
Gata6
Gata4
AP−2gamma
AP−2gamma
PGR
PSE
PR
1
2a
2b
3
4
5
6
Consensus Sequence Block
strand
−
+

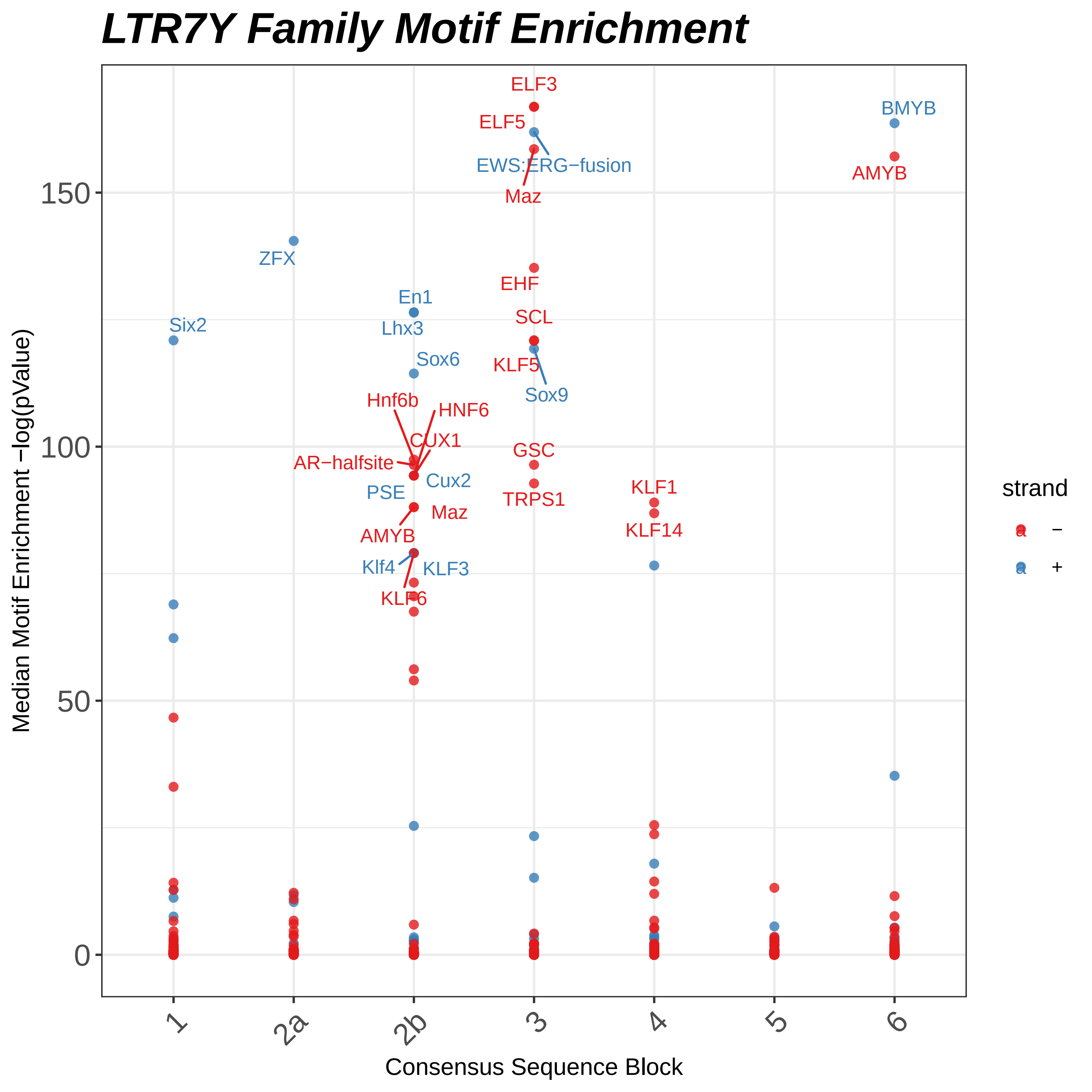

LTR7Y Family Motif Enrichment
Median Motif Enrichment −log(pValue)
Consensus Sequence Block
strand
−
+
0
50
100
150
1
2a
2b
3
4
5
6
ELF3
ELF5
EWS:ERG−fusion
Maz
BMYB
AMYB
ZFX
En1
Lhx3
EHF
SCL
Six2
Sox6
KLF5
Sox9
Hnf6b
HNF6
CUX1
GSC
AR−halfsite
Cux2
PSE
TRPS1
KLF1
Maz
AMYB
KLF14
Klf4
KLF3
KLF6
