## Supplemental figures for "Mosaic cis-regulatory evolution drives transcriptional partitioning of HERVH endogenous retrovirus in the human embryo"

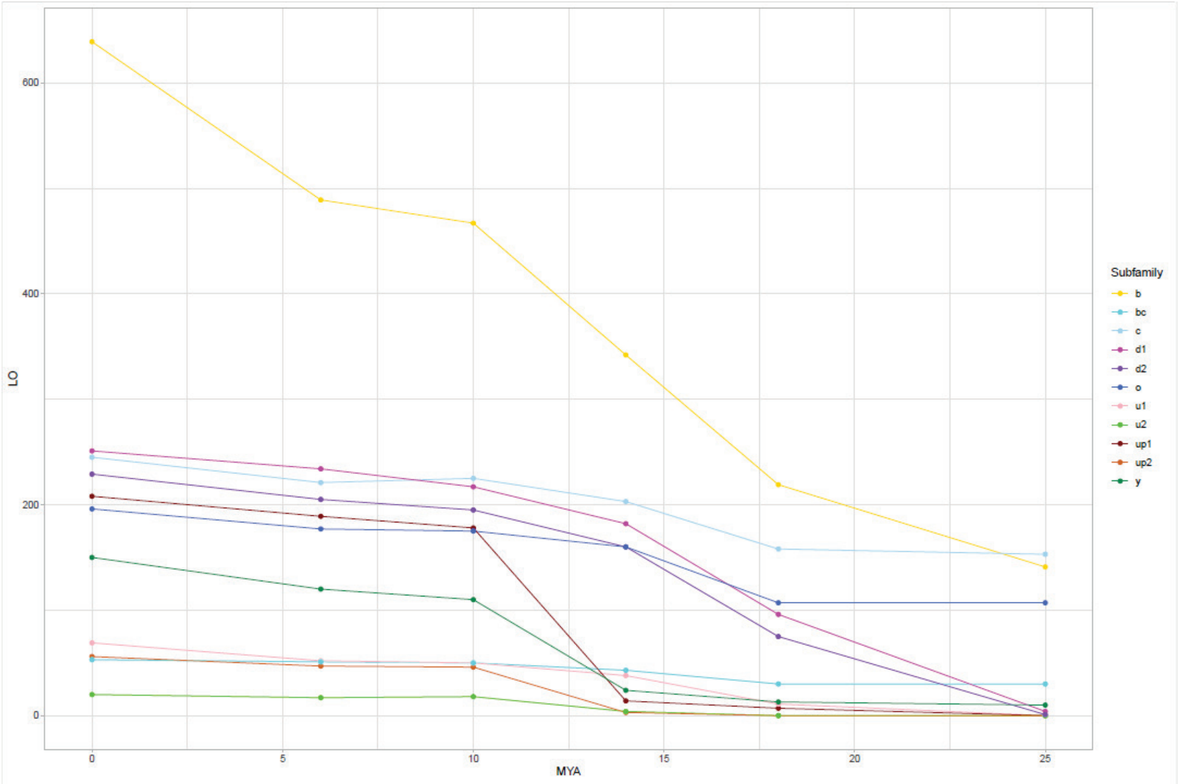


**Supplemental 1**: Gross reciprocal liftover from human to NHP. The absolute number of HERVH LTR that are in human are listed at MYA 0. Those that reciprocally liftover to chimpanzee, gorilla, orangutan, gibbon, and rhesus macaque are plotted as dots left to right with their relative divergence ages plotted as MYA (bottom). Points are connected with lines, where the slope approximates the rate of propagation of each subfamily between speciation events. This is a sister plot to main text Fig. 2a; the only difference is instead of proportion of reciprocal liftover, this is the gross number.

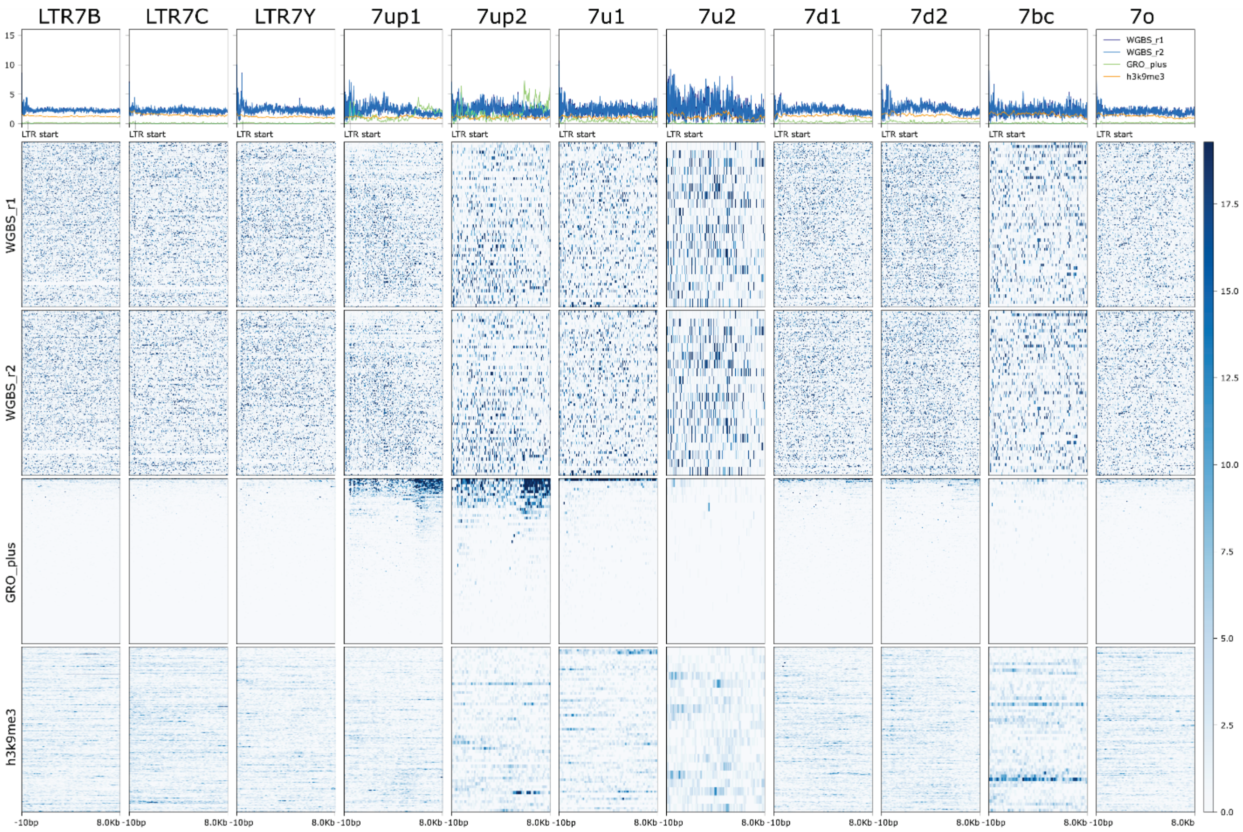


**Supplementary Figure 2**: Heatmap and aggregate signal of 2 replicates of whole-genome bisulfite sequencing (WGBS), GRO-seq (plus strand), and H3K9me3 in H1 cells. Visible window is 10bp upstream of the start of 5' (or solo) LTR and extends to 8kb after this point (~3kb past 3' LTR or ~7.5kb past solo into genomic DNA). Loci are sorted on GRO-seq data in the visible window.

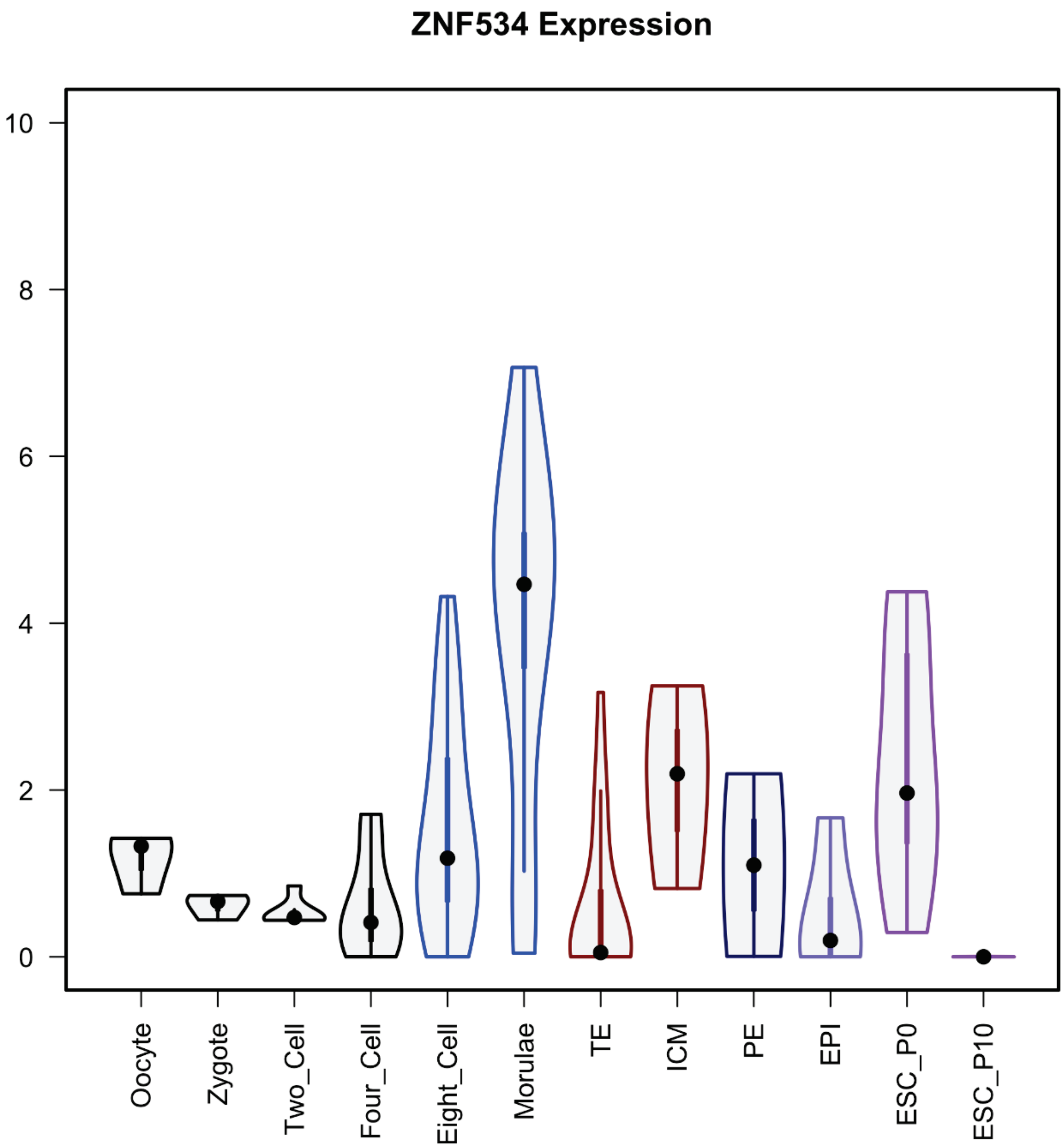


**Supplementary Figure 3:** Violin plots visualize the density and distribution of ZNF534 in early embryogenesis. Y-axis is [log TPM (transcripts per million)].

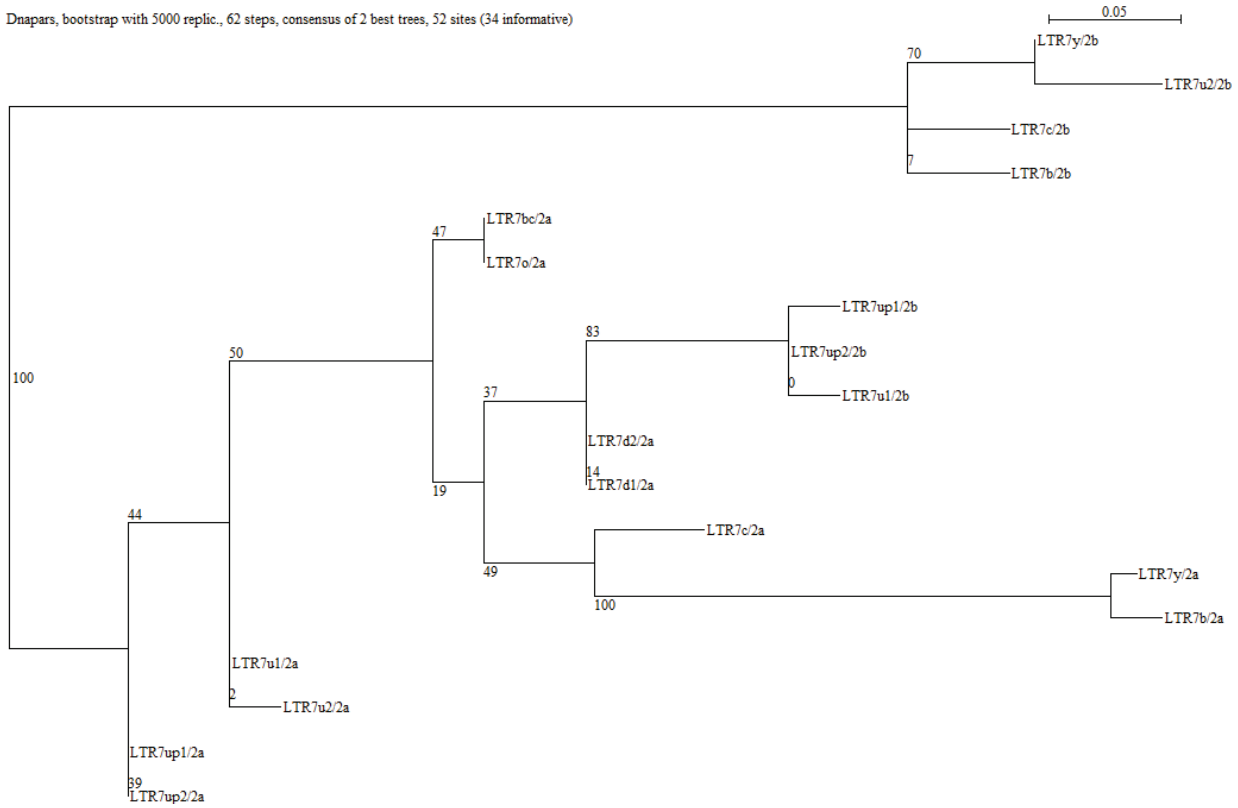


**Supplemental 4**: Parsimony tree of all consensus HERVH LTR blocks 2a and 2b. Sequence blocks 2a and 2b for all subfamilies were aligned using MUSCLE and the output was used to generate a parsimony tree in SEAVIEW (see methods).

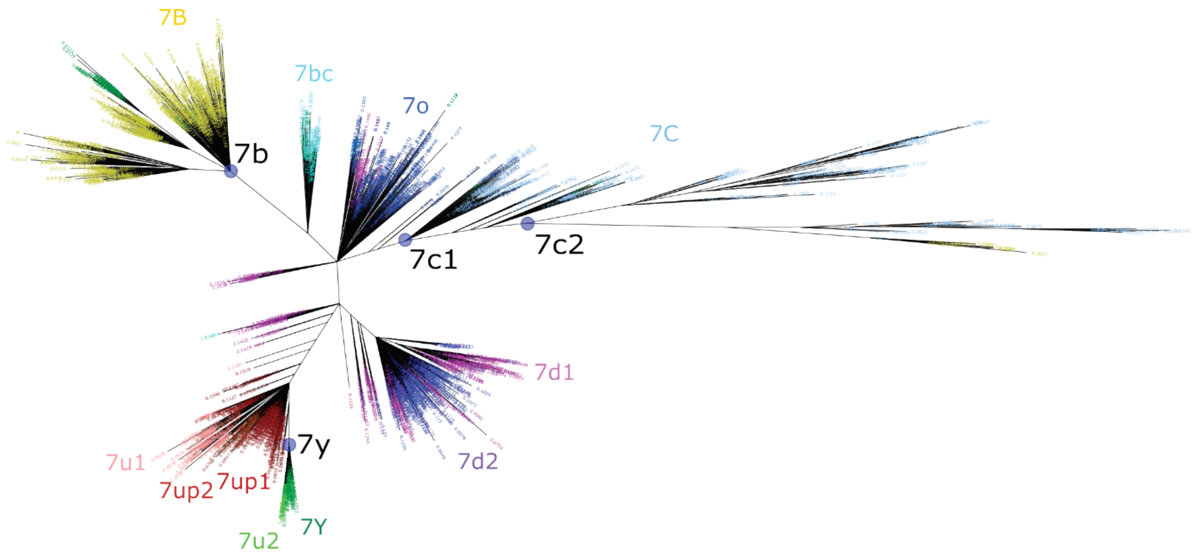


**Supplemental 5**: Majority-rule consensus sequences used for remasking of human genome. Consensus sequences for LTR7 subfamilies were generated from figure 1 (main text – see methods). Here, we show which sequences were included for the LTR7B, LTR7Y, and LTR7C (LTR7C1 + LTR7C2) consensus sequence generations (purple circles, black labels) using the tree from figure 1a. Previously annotated LTR7B/C/Y and LTR7 subfamily subdivision annotations from figure 1 (main text) are shown with colored node ages. The LTR7u2 sister clade was not included in the LTR7Y consensus generation.

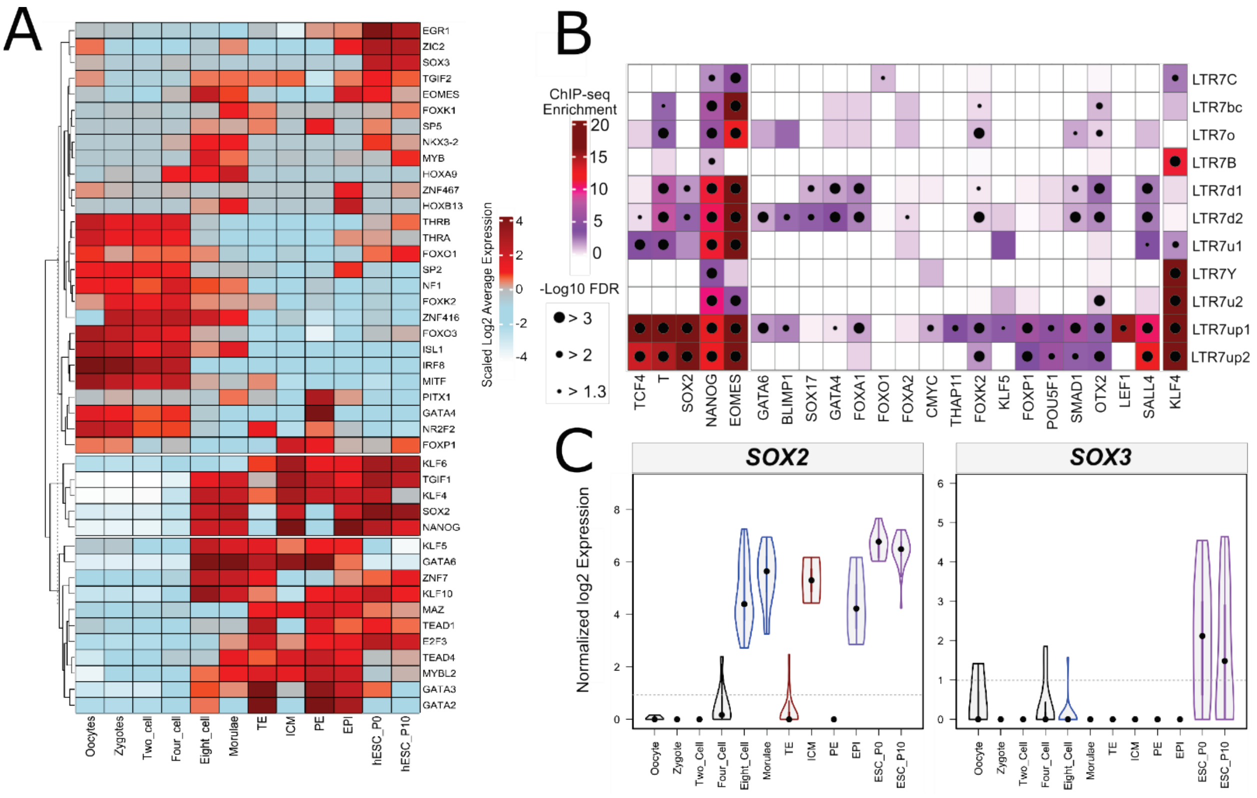


**Supplemental 6**: TF expression in the pre-implantation embryo and TF binding in embryonic stem cells. A) Heatmap showing averaged and scaled expression (log of normalized CPM values) of human transcription factors (TFs) with enriched motifs in LTR7-up1 subgroup and those are expressed ([log TPM (transcripts per million)] > 1) at least in one cell of early human embryos (Fig. supplement 8). Colour scheme is based on *z*-score distribution, from –4 (lightblue) to 4 (darkred). B) The merged dotplot and heatmap demonstrates overrepresentation analysis of pluripotency TFs binding in distinctive LTR7 subgroups. Color intensity corresponds to log scaled enrichment of the analyzed ChIP-seq peaks within LTR7 subgroups compared to randomized 500bps bins of human genome. Dots represent the significance (-log10 [FDR]) of ChIP-seq enrichment within a given LTR7 subgroup (see methods). C) Violin plots detail the spread of expression of SOX2/3 in the preimplantation embryo. Y axis is log2 TPM.

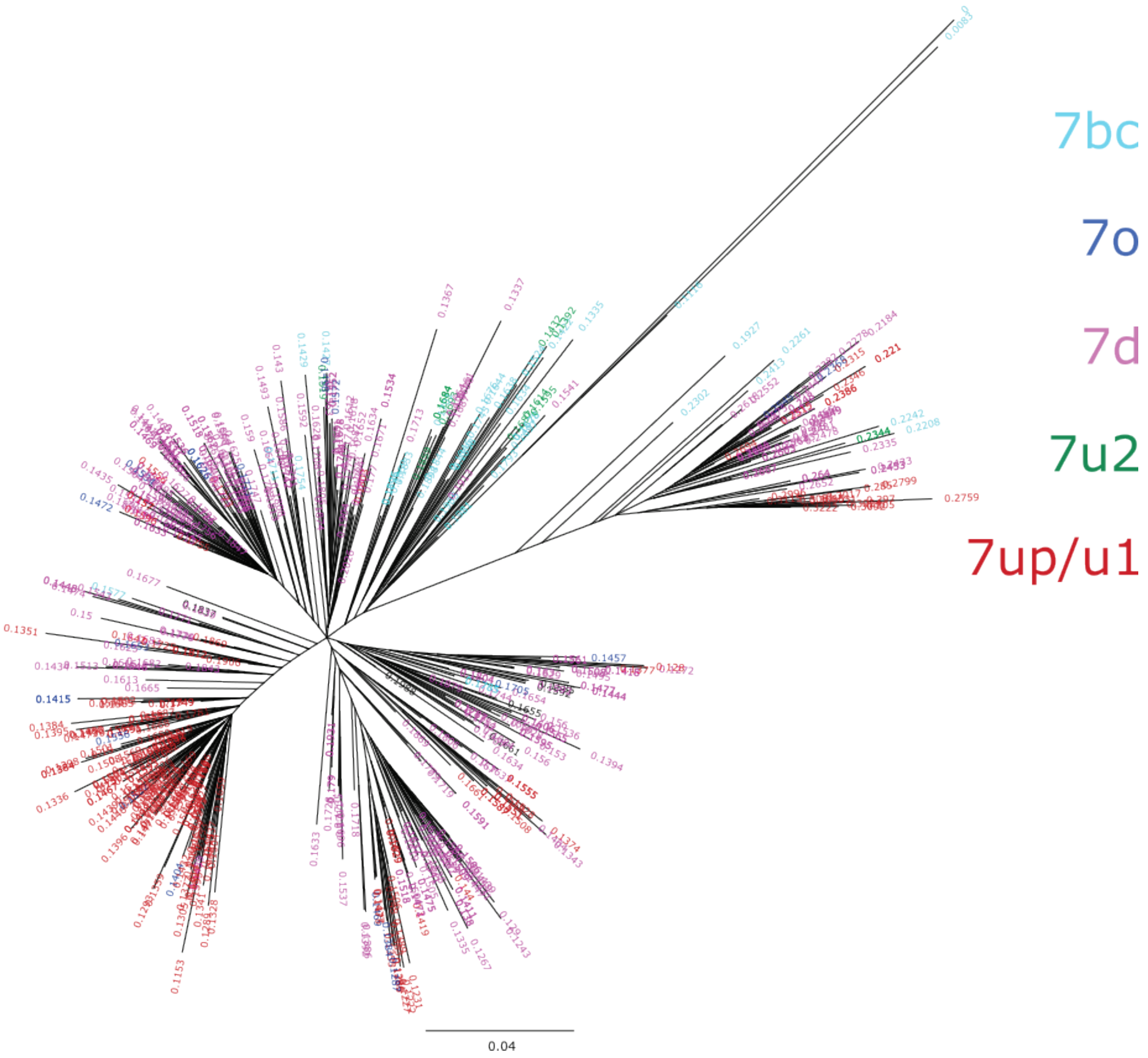


**Supplemental 7**: Phylogenetic tree from LTR7 reverse transcriptase domains (RVT). The consensus RVT domain was used to extract RT from individual LTR7 insertions. Relative node ages are shown at the end of the terminal branch lengths and are colored corresponding to their subfamily of origin. Only nodes with ultrafast bootstrap support >0.95 are shown. All displayed nodes are considered high confidence.

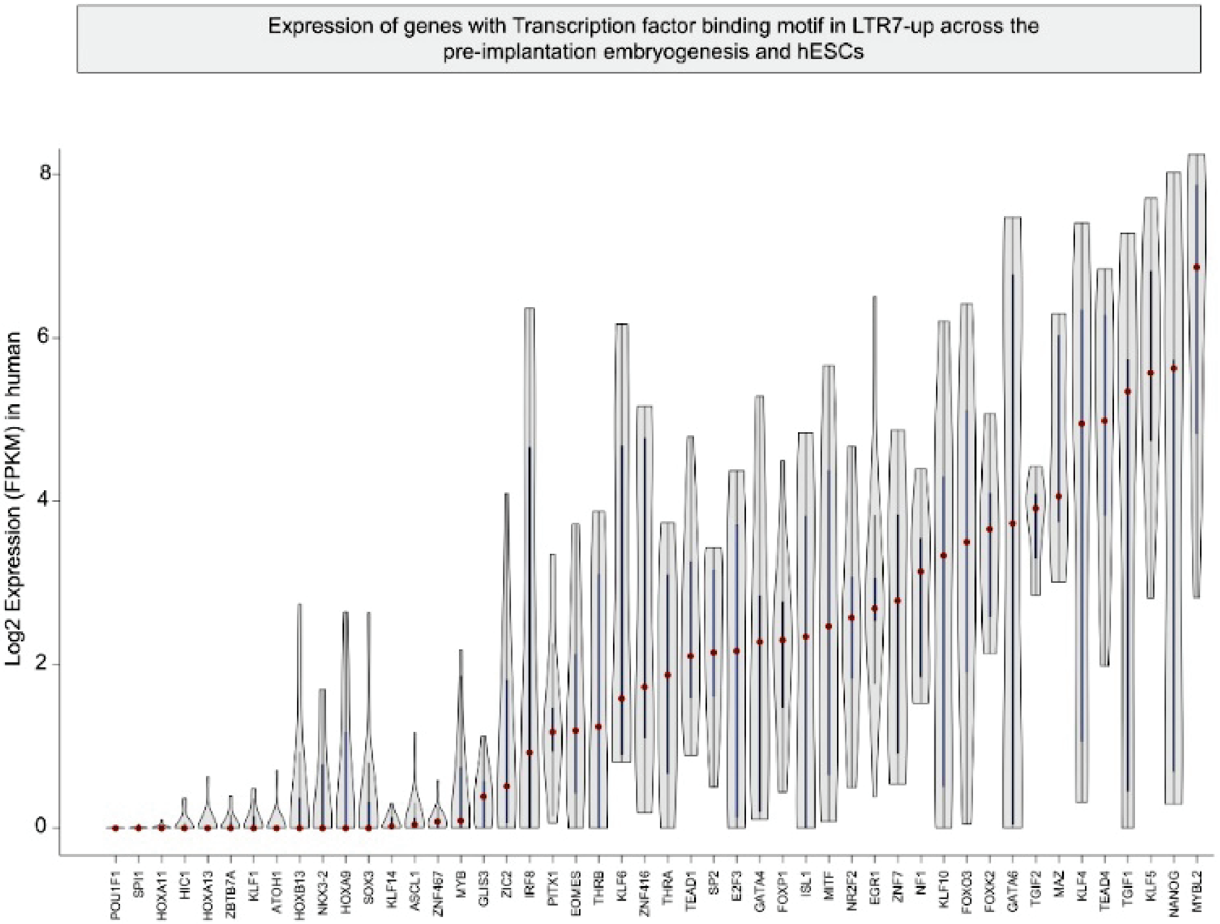


**Supplementary Figure 8**: Violin plots visualize the density and distribution of gene expression of TFs shown on supplemental 5 in analysed cells taken together from early human embryos and embryonic stem cells.
